# Widespread presence of direction-reversing neurons in the mouse visual system

**DOI:** 10.1101/826701

**Authors:** Yazan N. Billeh, Ramakrishnan Iyer, Iman A. Wahle, Shiella Caldejon, Séverine Durand, Peter A. Groblewski, Josh D Larkin, Jerome Lecoq, Ali Williford, Stefan Mihalas, Anton Arkhipov, Saskia E. J. de Vries

## Abstract

Direction selectivity – the preference of motion in one direction over the opposite – is a fundamental property of visual neurons across species. We find that a substantial proportion of direction selective neurons in the mouse visual system reverse their preferred direction of motion in response to drifting gratings at different spatiotemporal parameters. A spatiotemporally asymmetric filter model recapitulates our experimental observations.

---

Motion detection is a feature common to all visual animals, and recent work has shown strong parallels in these computations across many species^1–9^. Similar circuits in these animals underlie the comparison of visual information at two locations at two points in time, a computation that establishes direction selectivity wherein a neuron preferentially responds to motion in one direction over its opposite, or null, direction. Here, we report neurons in the mouse visual system that respond preferentially to motion in their null direction at different spatial and temporal frequencies.

Using data from the Allen Brain Observatory, a large-scale survey of visual responses in the mouse visual cortex recorded using 2-photon (2P) calcium imaging^10^, we analyzed responses to the drifting grating stimulus. This stimulus consisted of full-field sinusoidal gratings moving in 8 directions and at 5 temporal frequencies (TFs), but at a single spatial frequency (SF). Many neurons showed direction selectivity, and among these direction selective neurons we found some that reverse their direction preference in response to gratings at different TFs (**Fig. 1A**). We use the direction selectivity index (DSI, see Methods) to quantify the strength of direction selectivity at each TF, fixing the preferred and null directions to those determined at the preferred TF (TF_pref_, the TF evoking the largest mean response). A negative DSI thus indicates a reversal of direction preference (**Fig. 1B, C**). We term the neurons exhibiting this phenomenon Direction Reversing Neurons (DRNs).

**Figure 1:**
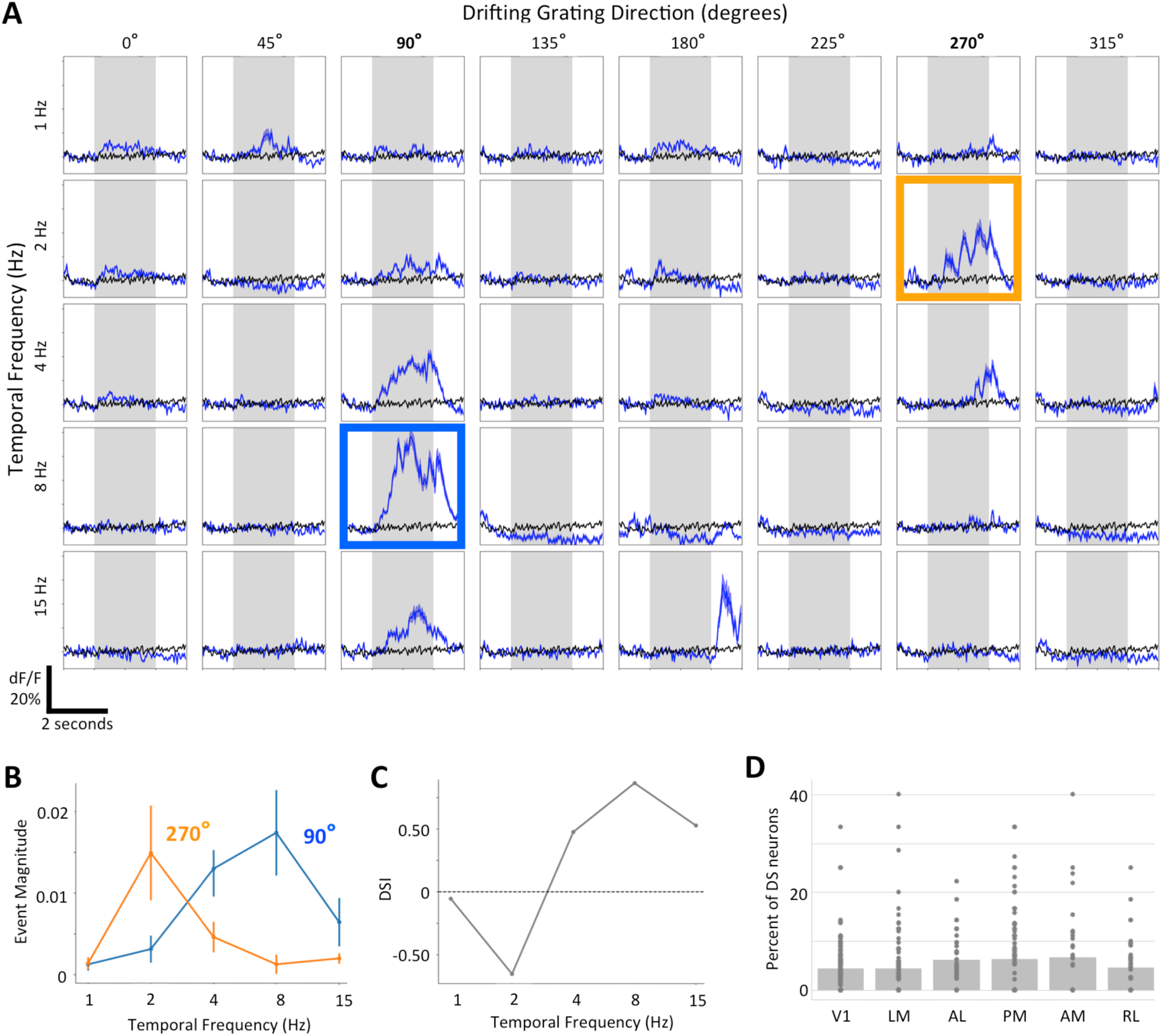
Direction Reversing Neurons (DRNs) prefer the null-direction as the stimulus temporal frequency (TF) changes. **A:** Example of a direction selective neuron’s calcium dF/F response to different TFs and directions of drifting gratings from the Allen Brain Observatory. Blue traces are the mean (± s.e.m.) response to the specified condition while the black traces are the mean (± s.e.m.) response to the blank-sweeps. The black trace is the same in each subplot. Note that the neuron’s strongest response is at a TF = 8 Hz and direction of 90°, which defines the *preferred condition*; the *null* direction is 270°. However, at TF = 2 Hz the neuron prefers the null direction of 270°. **B:** Responses (mean ± s.e.m.) of DRN in A at both the preferred and null direction as a function of TF. Note that hereafter our analysis employs events extracted previously^10^ from dF/F traces, rather than raw dF/F. **C:** Direction selectivity index (DSI) at each TF, where negative values indicate a preference for the null-direction. **D:** Percentage of direction selective neurons that are DRNs in each visual area. Each dot represents a single experiment, with the median for each area indicated by the bars. (n = 125, 101, 41, 87, 38, 40 experiments in V1, LM, AL, PM, AM, RL).

We imposed strict criteria to identify DRNs by comparing their responses at their preferred condition (the preferred direction at TF_pref_, blue box in **Fig. 1A**) and reversed condition (the TF at which the neuron has its largest mean response in the null direction, orange box in **Fig. 1A**). Candidate DRNs must have a DSI ≥ 0.3 for the preferred condition and exhibit a larger response to the null than preferred direction at the reversed condition. These neurons must also pass a strict bootstrap test (see Methods) to ensure the reversal is not a chance observation driven by a small number of outlier trials. 708 out of 12,515 direction selective neurons (6%) met these criteria. Furthermore, we observe DRNs in all visual areas recorded (**Fig. 1D**) and across all transgenically defined populations available in the dataset (**Fig. SI1**).

The presence of DRNs and the mechanism for this phenomenon have been explored in models of motion detection^8,9,11,12^ and even shown behaviorally in insects^1^. The fundamental ingredient is to have two spatially separated receptors, with one exhibiting a delayed response relative to the other, such as in the classic Hassenstein-Reichardt^1^ or Barlow-Levick^2^ models. Indeed, these models predict a reversal of direction preference due to aliasing in the neural response to periodic motion stimuli such as drifting gratings^8,9,11,12^. These mechanistic models make a testable prediction that direction selective neurons also reverse direction preference at different spatial frequencies (SFs). To test this prediction, we analyzed 2P experiments in which both the SF and the TF of the gratings were varied (see Methods). As predicted, we observed DRNs (using the same criteria as above) that reversed direction preference at different TF and/or SF conditions (**Fig. 2A, C**). This increased our DRN estimate to 15% of all direction selective neurons from 2P experiments (summed across all 6 cortical visual areas, 209/1411 neurons). Using an available dataset from extracellular electrophysiological recordings from V1 and the lateral geniculate nucleus (LGN)^13^ of the thalamus, we found a higher prevalence (21%, 40/193 neurons) of DRNs in V1 than in the 2P data (**Fig. 2B, C**), confirming that these reversals are not an artifact of calcium imaging and suggesting that the prevalence calculated from the Allen Brain Observatory data may be underestimated. Furthermore, finding DRNs in the LGN (29/139 neurons) demonstrates this is a common feature in the visual pathway.

**Figure 2:**
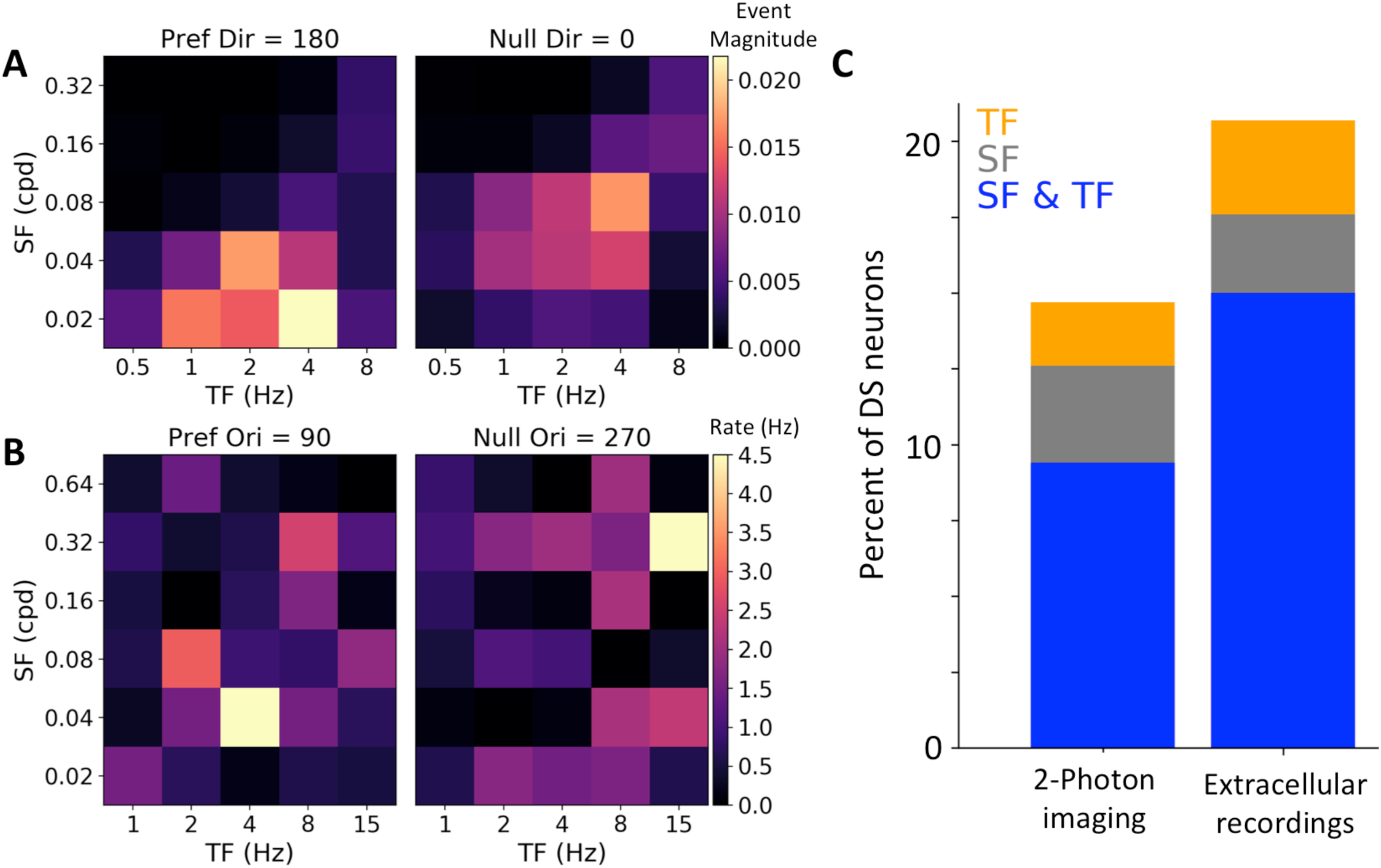
DRNs show reversal of the preferred direction due to changes in both TF and SF of the grating stimulus. **A:** Example DRN calcium response to drifting gratings of varying SFs and TFs. *Left:* Cell response in the SF and TF domain at its preferred direction. *Right:* Same cell’s response in the null-direction. Note that the cell responds to its preferred direction (180°) at SF = 0.02 cpd and TF = 4 Hz. However, the null-direction (0°) response is largest at SF = 0.08 cpd and TF = 4 Hz. **B:** Same as A for an example neuron from the electrophysiological dataset. **C:** Percentage of direction selective neurons that are DRNs for both recording modalities. Total direction selective neurons is 1411 neurons (2P) and 193 neurons (ephys).

These results show that direction preference reversals are a widespread phenomenon in the mouse corticothalamic visual pathway, constituting a fifth of direction selective neurons and occurring in every visual area we sampled. Therefore, we asked why these neurons were not reported more prominently in other mammalian systems. Part of the reason may have been that most studies didn’t have the dataset sizes available today to truly identify DRNs as a robust occurrence instead of chance observations. To the best of our knowledge, in mammalian studies, only one report showed neurons that reversed direction due to TF changes in areas 17 and 18 of the cat visual cortex^14^. The reversal occurred at very high temporal frequencies that are rarely studied. Given that the cat visual system was one of the most commonly studied for many decades, we sought to develop a model to explain the prevalence of DRN observations in mouse versus cat.

Recent experimental recordings in the mouse demonstrated a spatiotemporal asymmetry in the convergence patterns of LGN projections onto Layer 4 V1 neurons, which was essential in establishing direction selectivity in these neurons^15^. Unlike the classic models that incorporated a time delay between receptors, the mechanism here involves different temporal response profiles between two input components: a sustained, long lasting component, and a transient, short lasting component (similar to the OFF pathway motion detection system in the fly T5 neurons^16,17^). Based on this, we developed a model that similarly splits a neuron’s receptive field into two subfields (transient and sustained), parameterized from either mouse or cat measurements.

Our model consisted of simple point-receptor subfields with square-wave temporal responses (**Fig. 3A, B**). We first characterized direction selectivity as a function of the TF and SF of the drifting grating stimulus using mouse parameters^18,19^. We find that the relative phase of the subfield responses can shift from completely out-of-phase to completely in-phase depending on both the direction and SF of the stimulus (**Fig. 3C**). This phase shift causes reversal in the preferred direction as the SF varies (**Fig. 3D**). Moreover, this model exhibits direction reversals due to changes in TF (**Fig. 3E**) as in previous models^1,2,8,9,15–17^. Note that a difference in time-constants between the two subfields is necessary for direction selectivity to emerge at all (see heat-maps in **Figs. 3D, E**). Finally, while the model used here has an ON sustained and OFF transient subfields, the observed results hold for any other ON/OFF combinations, with the only requirements being spatial offset and temporal asymmetry (**Fig. SI2**). Indeed, the spatiotemporal asymmetry of subfields is sufficient for establishing direction selectivity and direction reversals.

**Figure 3:**
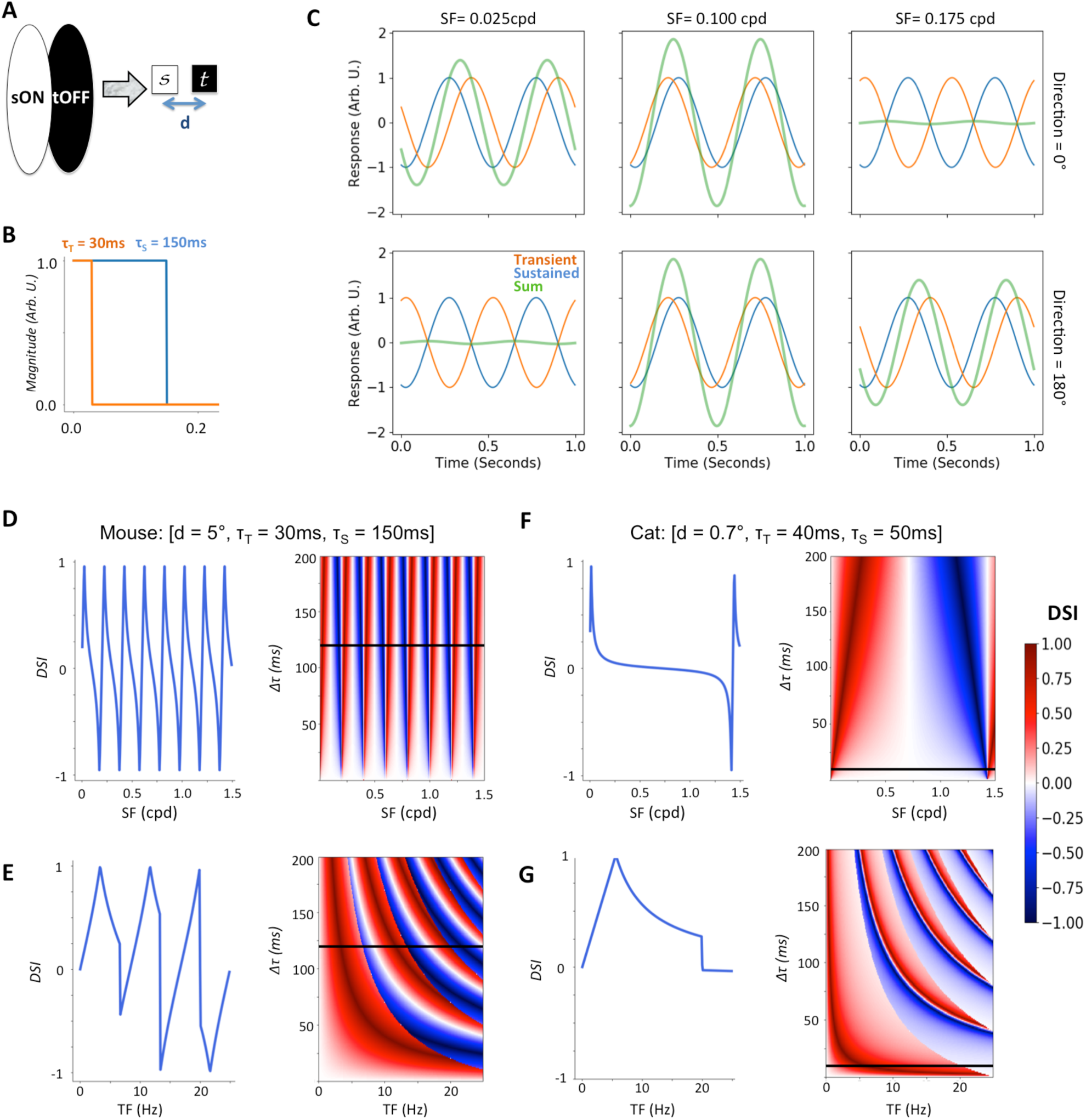
Model of a basic spatiotemporal asymmetry in receptive subfields predicts DRN responses. **A:** Schematic of a simple cell with spatially separated opponent subfields, here with different temporal parameters. This model can be reduced to two point receptors separated by a distance *d* in the visual space. **B:** Temporal filters of the point receptors used (simple square waves). The response amplitude was +1 for ON filters (shown) and −1 for OFF filters. The time constants *τ*_*T*_ and *τ*_*S*_ determine the duration of the responses. **C:** Subfield responses for the model at different drifting grating SFs for two directions (0° and 180°). The plots use mouse parameters (see D) and the drifting grating TF is kept constant at 2 Hz. *Left:* Responses at SF= 0.025 cpd showing near identical phase responses at 0°and antiphase responses at 180°, resulting in a strong preference for motion at 0°. *Middle:* Responses at SF= 0.1 cpd exhibiting equal phase offsets in both directions. *Right:* Responses at SF= 0.175 cpd, showing antiphase responses at 0° and near identical phase responses at 180°, resulting in a strong preference for motion at 180°. **D:** DSI as a function of SF for the “mouse” model. *Left:* plot of DSI against SF for a *Δτ* = 120 *ms* (*Δτ* = *τ*_*S*_ − *τ*_*T*_). *Right:* heatmap of DSI as a function of SF and *Δτ*. The black line corresponds to the slice represented on the left. **E:** Same as (D) but for the TF. **F:** Same as (D), but for the “cat” model. Direction reversal only occurs at very high SF values that are rarely tested. **G:** Same as (E), but for the “cat” model. Reversals occur only at high TFs. The default drifting gratings parameters are TF = 2Hz and SF = 0.04 cpd for this figure.

We then employed the same model with parameters derived from recordings in the cat^9^. The major difference from the mouse is that the cat visual system exhibits a substantially higher acuity (i.e. a smaller distance between subfields) and faster responses for both the transient and sustained components. Consequently, the “cat” model predicts reversal of direction preference at much higher SF/TF values that are near the physiological response limits of cat visual cortex neurons (**Fig. 3F, G**)^20,21^. This agrees with the previous study that reported direction reversal in cat visual neurons at high TFs^14^. These results suggest that DRNs have not been widely observed in mammals because they are truly rare in the cat (and presumably in other species possessing high visual acuity, such as primates) whereas mouse studies until recently did not sample enough neurons to clearly identify that DRNs are not a chance occurrence.

It is possible that our results underestimate the true prevalence of DRNs in the mouse. The fact that our model predicts multiple reversals within physiological limits (**Fig. 3D, E**) implies that more DRNs could be revealed if probed with a larger range or finer sampling of SF and/or TF. This raises the question of whether all mouse direction selective neurons would show direction reversals in response to a periodic stimulus if probed with the right conditions. Previous studies in the fly have shown that under certain conditions, such as a large overlap between subfields or temporal filters with low pass filter properties, the prediction of direction reversal is eliminated^11^. Given that many direction selective neurons in the mouse visual system are not described by well-separated subfields, this could account for non-DRN direction selective neurons. It is also possible that some direction selective neurons receive lateral or feedback connections that counteract the reversing signals, or that reversals happen under conditions where responses are too small and indistinguishable from noise. We thus expect that DRN abundance in the mouse may be higher than what we observed, but at the same time it is likely that not every direction selective neuron is a DRN.

In this work, we have demonstrated the surprising prevalence of DRNs throughout the mouse visual system. It remains to be seen whether these neurons are advantageous in visual processing, or whether they are an epiphenomenon elicited by periodic motion stimuli. The latter could explain perceptual phenomena such as illusory motion reversals, including the “wagon wheel” illusion^22,23^. In either case, our results suggest that the difference in receptive field sizes between mice and carnivores, such as cats, or primates impacts more than just spatial acuity, but also other aspects of visual processing including motion detection.

## Acknowledgements

We thank the Allen Institute founder, Paul G. Allen, for his vision, encouragement, and support.

## Methods

### Datasets

We analyzed data from the Allen Brain Observatory^10^. This dataset consists of neural activity recorded using 2P calcium imaging of transgenically expressed GCaMP6 in a number of different transgenic lines in 6 different cortical areas. We limited our analysis to the GCaMP6f data in the dataset. We studied the responses to the drifting grating stimulus, which consisted of a full field drifting sinusoidal grating that was presented at five temporal frequencies (1, 2, 4, 8, 15 Hz) and eight different directions (from 0° to 315°, 45° steps between each), and at spatial frequency of 0.04 cpd. Each grating was presented for 2 seconds, followed by 1 second of mean luminance gray before the next grating. Each grating was presented 15 times in a random order, with blank-sweeps interleaved roughly every 20 trials.

The calcium data were processed in a standardized manner, and all analysis here was performed using events extracted from the ΔF/F traces^10^.

### Spatial and temporal frequency 2P experiments

Data was collected using the same data collection pipeline as the Allen Brain Observatory^10^ and processed using the same image processing and event detection methods. The stimulus consisted of a full field drifting sinusoidal grating that was presented at five spatial frequencies (0.02, 0.04, 0.08, 0.16, 0.32 cpd), five temporal frequencies (0.5, 1, 2, 4, 8 Hz), and 4 directions (0, 90, 180, 270°). Each grating was presented for 2 seconds, followed by 1 second of mean luminance gray before the next grating. Each grating condition (direction, SF, TF combination) was presented 15 times. Trials were randomized with blank sweeps interleaved approximately once every 80 trials.

Data was collected from Cux2-CreERT2;Camk2a-tTA;Ai93 mice imaged at 175 μm below the cortical surface in layer 2/3, across visual areas V1, LM, AL, PM, AM and RL. Within V1, data was also collected from Rorb-IRES2-Cre;Camk2a-tTA;Ai93 imaged at 275 μm below the cortical surface in layer 4, Rbp4-Cre_KL100;Camk2a-tTA;Ai93 imaged at 375 μm below the cortical surface in layer 5, and Ntsr1-Cre_GN220;Ai148 imaged at 550 μm below the cortical surface in layer 6. (These Cre lines and imaging depths match those used in the Allen Brain Observatory. See de Vries, Lecoq, Buice *et. al* 2019^10^, for further Cre line and imaging details. In total 3,466 neurons were imaged in 77 experiments.

### Extracellular electrophysiology dataset

We analyzed previously published extracellular electrophysiological dataset^13^. We used data from V1 and LGN recorded in awake mice. The stimulus consisted of a full field drifting sinusoidal grating that was presented at six spatial frequencies (0.02, 0.04, 0.08, 0.16, 0.32, 0.64 cpd), five temporal frequencies (1, 2, 4, 8, 15 Hz), and 8 directions (45° steps). Each grating was presented for 3 seconds, followed by 1 second of mean luminance gray before the next grating. Each grating condition (direction, SF, TF combination) was presented at least 7 times. Trials were randomized with blank sweeps interleaved. In total 1,311 neurons were recorded in 14 mice.

### Analysis

The preferred condition of a neuron is defined as the direction, TF (and SF, where applicable) that elicited the largest mean response across trials (blue box for the example neuron in **Fig. 1A**).

The direction selectivity index (DSI) was computed using extracted events or, for electrophysiological recordings, from spikes, by taking the ratio of the difference between the preferred and null response relative to their sum:

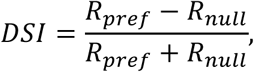

where *R*_*pref*_ and *R*_*null*_ are the mean event magnitude (2P) or mean spike count (electrophysiology) over the duration of the corresponding grating, averaged over all trials, for the preferred and null directions, respectively. The DSI was computed at each TF (and SF, where applicable) using the preferred and null directions defined at the preferred TF (or preferred TF/SF combination). Negative DSI values thus indicate a stronger response in the null direction than the preferred.

### DRN selection via Bootstrap

To ensure that DRN identification was not driven by outlier response trials, a bootstrapping process was used to generate additional response trials from the distribution approximated by the data. For a given neuron, a 15-trial sample was drawn randomly with replacement from the 15 responses to the preferred and null conditions, respectively. This procedure of resampling responses to create equal sized datasets was done 1000 times for every neuron. A sample with 1) a DSI ≥ 0.3 and 2) a larger mean response to the null than preferred direction at the reversed condition was considered a positive sample. Candidate DRNs with greater than 95% positive samples out of 1000 generated samples were identified as DRNs.

### Spatiotemporal asymmetric receptive field model

The model we used to approximate integration of spatiotemporally asymmetric inputs by a neuron consisted of two point receptors separated by a distance in the visual space, *d*. In addition, the second key component for attaining direction selectivity in our model was that both receptors had different time constants [Lein and Scanziani 2018]. We termed the slower filter the sustained filter (*F*_*S*_) and the faster filter the transient filter (*F*_*T*_). Again in the interest of simplicity we used standard square wave functions (**Fig. 2A**, code available at github.com/saskiad/direction_flipping/tree/master/model_code) with ON filters being described by:

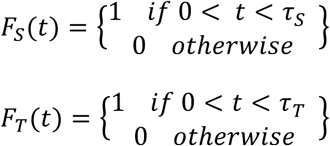

where *τ*_*S*_ and *τ*_*T*_ are the time-constants (duration) of the sustained and transient receptor units, respectively. The OFF filters simply had an amplitude of −1.

For simplicity, one can consider the case of sustained and transient receptors with the line connecting them at zero degrees (as illustrated in **Fig. 2A**). Also the sustained receptor is defined as the origin and the transient receptor at distance *d* away:

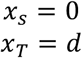

If a sinusoidal drifting grating (assumed at full contrast), with spatial frequency *k* and temporal frequency *ω* = 2*πf*, is presented, the input on the receptor units can be described as:

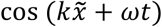

where *t* is the time from stimulus onset. Here 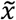 is the position of the receptor after adjusting for the orientation of the drifting grating input. Since we are considering the receptors are on the zero degree (horizontal) line:

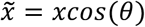

where *θ* is the drifting grating’s direction. This is simply the projection of the grating on the horizontal axis represented by *x*. For example, consider when the drifting grating is moving at 90° (vertically); in that case both receptors will always be in the exact same phase (“seeing the same thing”) and hence the spatial component is irrelevant. If the drifting grating is moving at 0°, however, then the position of the receptor (*x*) and the spatial frequency (*k*) account for the phase difference between the receptors. As such, the input stimulus to the sustained receptor can be expressed as:

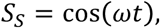

since we consider it to be at the origin. Similarly, the input stimulus to the transient receptor can be expressed as:

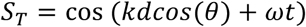

Thus the responses of the filters can be described by the convolution of the filter with the stimulus (recall the square-wave profile of *F*_*S*_ and *F*_*T*_):

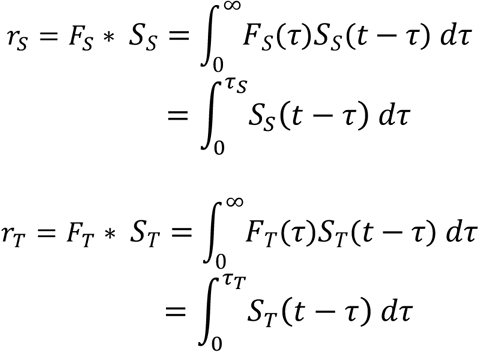

where *τ*_*S*_ and *τ*_*T*_ are the time-constants (duration) of the sustained and transient receptor units, respectively, that only impacted the integral limits (since unitary square waves filters are modeled).

Given that the responses are variable in magnitude due to the input frequency of the grating, we normalized the responses before adding both components. Moreover, the total response would, as expected, oscillate at the stimulus frequency and we thus defined the net response at a specific orientation as the maximal value reached:

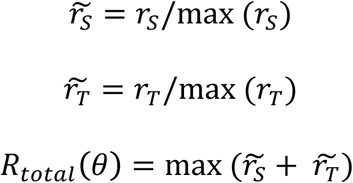

Similar to the experimental data, we finally calculated the DSI by taking the difference between the preferred (0°) and null (180°) directions and dividing by their sum:

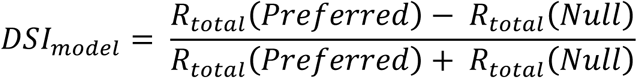

## Contributions

Conceptualization: YNB, SEJdV. Methodology: YNB, SEJdV, RI, AA, SM. Formal Analysis: YNB, SEJdV, RI, IW. Data Collection: JL, JDL, SC, AW, PG, SD; Writing-Original: YNB, SEJdV; Writing – Review and Editing: YNB, SEJdV, RI, AA.

## Supplemental Information

**Figure S1:**
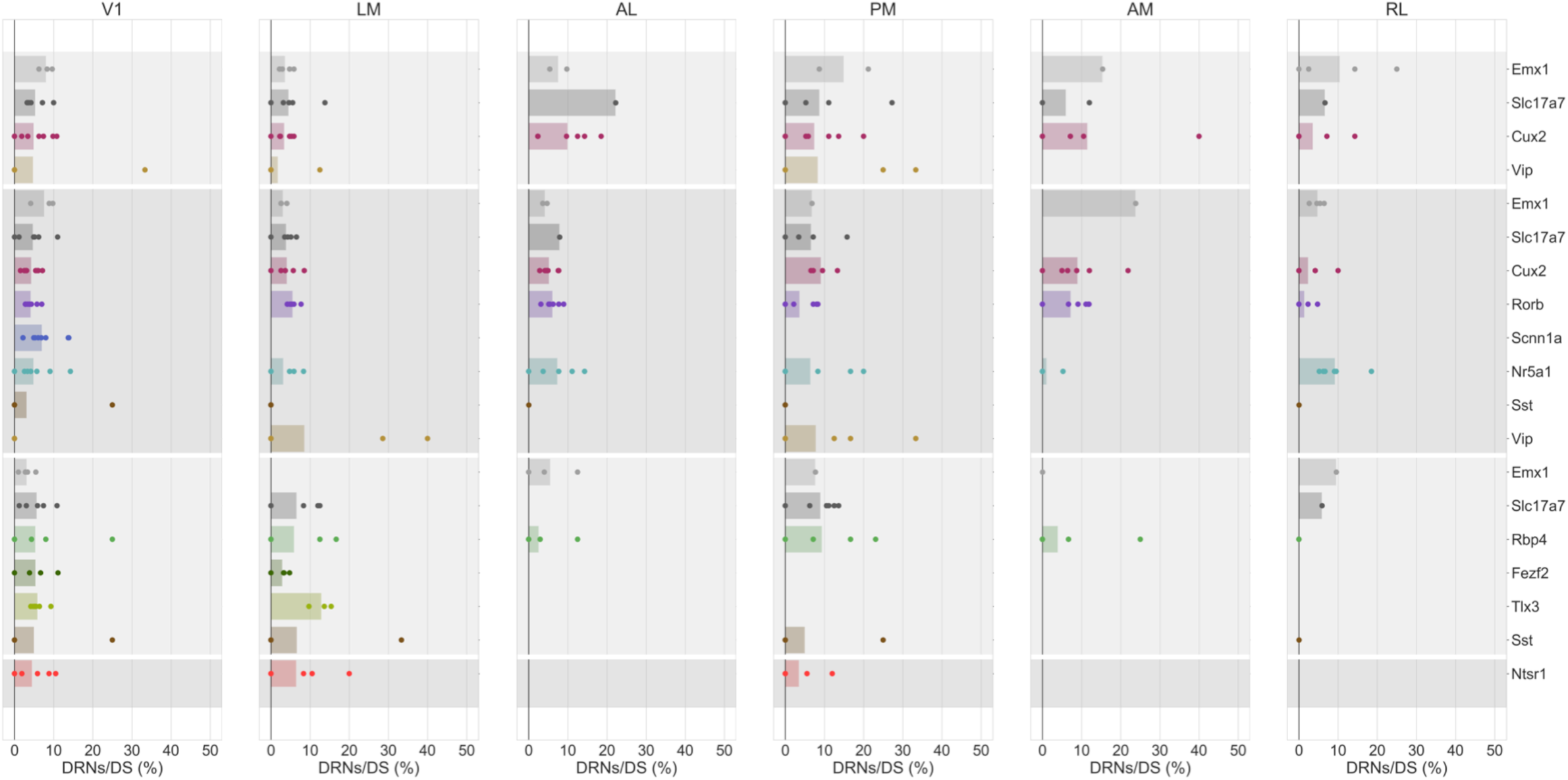
Strip plot of percent of DS neurons that are DRNs for each Cre line and area. Dots represent individual experiments and the bar represents the median across experiments.

**Figure S2:**
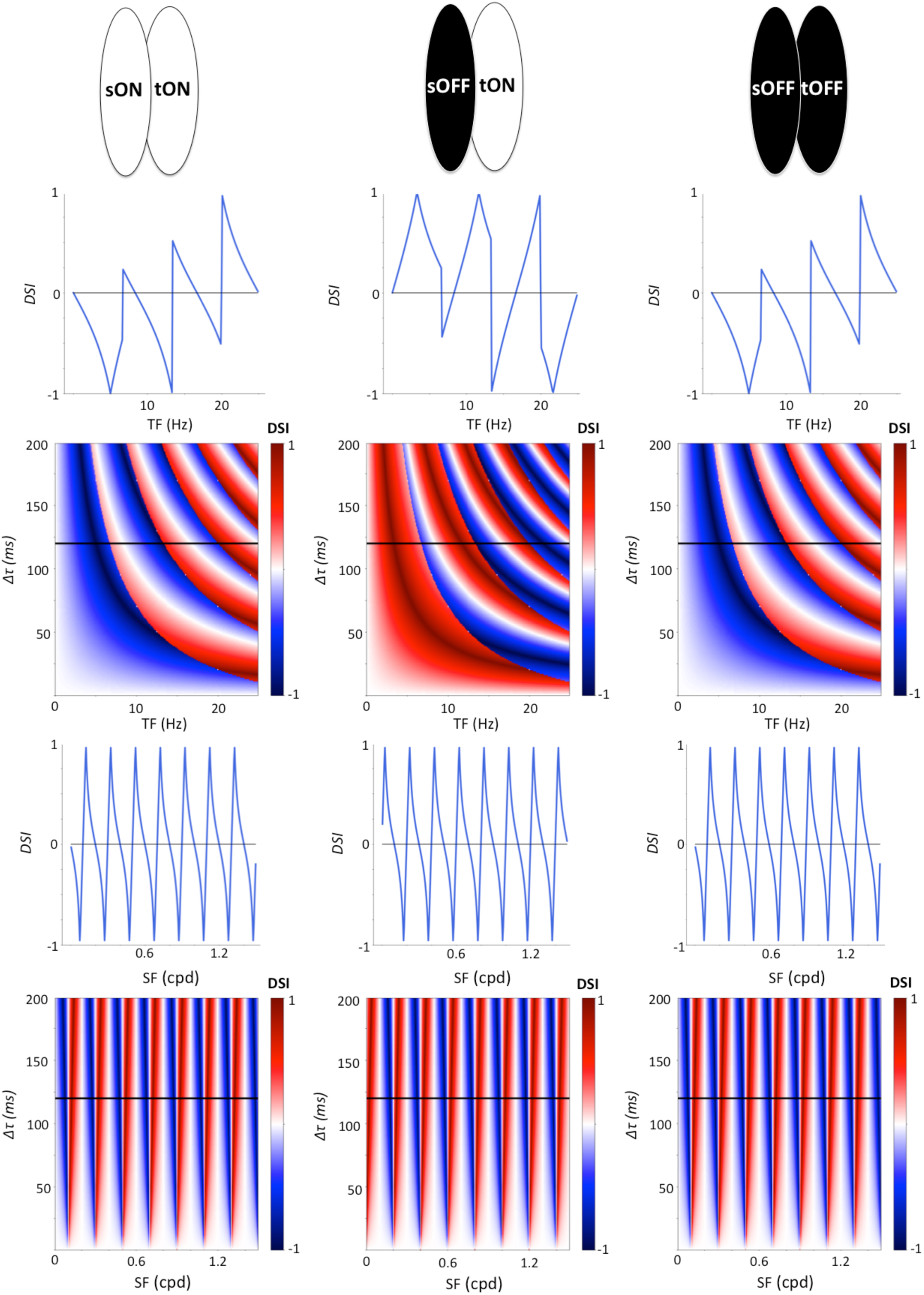
Spatiotemporal asymmetric model predicts reversal regardless if units are ON or OFF. All parameters use the “mouse” model. *Top*: Schematic spatially separated opponent subfields for every column in the figure. *Second row:* DSI as a function of TF for the “mouse” model for a *Δτ* = 120 *ms* (*Δτ* = *τ*_*S*_ − *τ*_*T*_). *Third Row:* heatmap of DSI as a function of TF and *Δτ*. The black line corresponds to the slice represented on the second row. *Fourth row:* Same as second row but for the SF. *Fifth row:* Same as third row but for SF. The default drifting gratings parameters are TF = 2Hz and SF = 0.04 cpd for this figure.

